# Inhibition of 3-mercaptopyruvate sulfurtransferase enhances CD8⁺ T-cell antitumor immunity

**DOI:** 10.1101/2025.09.22.675323

**Authors:** Muriel Urwyler, Marietta Margareta Korsos, Eloise Dupuychaffray, Kelly Ascenção, Maria Petrosino, Karim Zuhra, Jérémie Tachet, Pragallabh Purwar, Montserrat Alvarez, Aurélien Pommier, Mohamed Z. Gad, Reham M. Abdel-Kader, Csaba Szabo, Carole Bourquin

## Abstract

Hydrogen sulfide (H₂S) is a redox-active gasotransmitter implicated in tumor progression and immune regulation. The enzyme 3-mercaptopyruvate sulfurtransferase (3-MST) is a key contributor to endogenous H₂S and polysulfide production, but its role in tumor–immune interactions remains poorly defined. Here, we show that 3-MST is the most abundantly expressed H₂S-synthesizing enzyme in human renal cell carcinoma cells (RCC) and that high 3-MST expression correlates with reduced patient survival. Pharmacological inhibition of 3-MST lowered intracellular H₂S levels in Renca renal carcinoma cells, suppressed proliferation, induced apoptosis, and increased surface expression of the immunogenic markers CD70, CD86, and PD-L1. In immune cells, partial inhibition of 3-MST promoted T cell activation, as evidenced by increased CD69 expression on both CD4⁺ helper and CD8⁺ cytotoxic T cells. In contrast, complete inhibition of 3-MST, achieved by high concentrations of the inhibitor, modestly reduced CD8⁺ T cell proliferation. Functionally, 3-MST inhibition potentiated antigen-specific CD8⁺ T cell-mediated killing of tumor cells, an effect further amplified by PD-L1 blockade. These results establish 3-MST as a redox-sensitive metabolic driver of tumor growth and immune evasion in RCC and demonstrate that its inhibition can boost antitumor immune responses, offering a potential avenue for combination immunotherapy.

## Introduction

Renal cell carcinoma (RCC) accounts for approximately 90% of all kidney cancers and remains a significant clinical challenge due to its resistance to conventional therapies [1,2]. Immunotherapy represents an important option for patients with RCC and is often used in combination with conventional treatments. The mainstay of immunotherapy in RCC are immune checkpoint inhibitors, which prevent immunosuppression of CD8^+^ T cells and restore their anticancer functionality [1]. Checkpoint inhibition has improved outcomes in many patients with advanced RCC, but therapeutic resistance and limited response rates highlight the need for novel strategies to sensitize these tumors to immune-mediated elimination [3].

Hydrogen sulfide (H₂S) is an endogenously produced gaseous signaling molecule involved in a wide range of physiological and pathophysiological processes, including mitochondrial respiration, redox balance, and inflammation [4]. In the context of cancer, H₂S has been implicated in supporting tumor growth and survival by promoting metabolic flexibility, angiogenesis, and resistance to oxidative stress [5–8]. Beyond its effects on tumor cells, H₂S also influences the immune response by regulating T cell activation, cytokine production, and the expression of immune checkpoint molecules and may thus impact antitumor immunity [9,10]. In mammalian cells H₂S is synthesized by three main enzymes: cystathionine β-synthase (CBS), cystathionine γ-lyase (CTH, also known as CSE), and 3-mercaptopyruvate sulfurtransferase (3-MST, encoded by *MPST*) [4]. While these enzymes have been extensively studied in cancer metabolism and inflammation [11–13], their precise roles — and that of 3-MST in particular — in antitumor immunity remain insufficiently characterized [14].

3-MST is partially localized to the mitochondria [15], where it catalyzes the conversion of 3-mercaptopyruvate into H₂S and other reactive sulfur species, thereby contributing to redox buffering and cellular antioxidant defenses [16,17]. Recent studies suggest that 3-MST contributes to the metabolic adaptability and survival of tumor cells, particularly under stress conditions such as hypoxia or oxidative challenge. However, its role in tumor immune evasion or immune modulation has not been investigated [18]. Interestingly, 3-MST activity is detected in many tissues, but the highest activity of 3-MST was measured in the kidney [14].

In this study, we examined the role of 3-MST in tumor metabolism, immunogenicity, and immune cell interactions using the murine Renca renal carcinoma model. We used 2-[(4-hydroxy-6-methylpyrimidin-2-yl)sulfanyl]-1-(naphthalen-1-yl)ethan-1 (HMPSNE), a selective pharmacological inhibitor of 3-MST, to assess how suppression of this enzyme affects tumor cell growth and CD8⁺ T cell-mediated antitumor responses. Our findings demonstrate that 3-MST inhibition impairs tumor cell viability, enhances the expression of immunoregulatory surface molecules, and promotes antigen-specific cytotoxic T cell activity, particularly when combined with immune checkpoint inhibition. These results highlight 3-MST as a potential target for modulating tumor-immune interactions and improving immunotherapeutic strategies in patients with renal cancer.

## Methods

### Reagents

6-Methyl-2-((2-(naphthalen-1-yl)-2-oxoethyl)thio)pyrimidin-4(3H)-one (I3MT-3, HMPSNE, Otava Chemicals, CAS 459420-09-8) was first prepared as a 500 mM stock solution in DMSO, then further diluted 1:10 in DMSO. From this intermediate solution, a 5 mM working solution of HMPSNE was prepared in complete cell culture medium for experimental use.

### Mice

Female C57BL/6J and OT-I (C57BL/6-Tg(TcraTcrb)1100Mjb/Crl) mice aged 6–16 weeks were used as cell donors. Animals were maintained under specific pathogen-free conditions at the Centre Medical Universitaire, University of Geneva in accordance with Swiss animal welfare regulations. Mice were obtained from Charles River Laboratories (Saint Germain Nuelles, France).

### Cell lines

The Renca cell line (ATCC, CRL-2947) was genetically modified to express MHC Class I H2-K^b^ instead of the native H2-K^d^ and to express GFP (Renca H2-K^b^ GFP), as described previously [19]. Cells were cultured in RPMI-1640 medium (Gibco, #21875034) supplemented with 10% FBS (Capricorn Scientific, Germany), 100 U/mL penicillin/streptomycin (Gibco, #15140122), 1% non-essential amino acids (Gibco, #11140035), 1 mM sodium pyruvate (Gibco, #11360039), and 2 mM L-glutamine (Gibco, #25030024) (complete RPMI medium) at 37 °C in a 5% CO₂ humidified atmosphere. Cells were seeded in 96-well plates (Greiner CellStar, #655180) and allowed to adhere for 24 h prior to HMPSNE treatment. Tumor cell proliferation was monitored by GFP intensity using the Incucyte® Live-Cell Analysis System (Sartorius). Data were expressed as GFP signal normalized to the starting point (T₀ = 1) for each well.

### Detection of H_2_S

H_2_S generated in living cells was measured using the AzMC probe as previously described [20,21]. Cells were seeded in sterile black 96-well plate with optical bottom at 3000 cells/well density in 100 µl of complete culture medium and incubated over-night at 37°C and 5% CO_2_. The day after cells were treated with different concentrations (3, 10, 30 and 100 µM) of HMPSNE. After 48h, culture medium was replaced with HBSS buffer supplemented with 1 mM 7-azido-4-methylcoumarin (AzMC) H_2_S-sensitive probe and D-glucose 4.5 g/l and further incubated for 2 h. Dye’s specific fluorescence was visualized using a Leica DFC360 FX microscope and images were captured with Leica Application Suite X software (Leica Biosystems Nussloch GmbH, Germany). Images were analyzed with ImageJ software (v. 1.8.0; NIH, Bethesda, Maryland, USA) and data graphed with GraphPad Prism 8 (GraphPad Software Inc.; San Diego, California, USA).

### Mouse splenocytes

To generate single cell suspensions, mouse spleens were passed through a 40 µM filter and treated with Pharm Lyse™ red blood cell lysis buffer (BD, #555899). CD8a^+^ T cells were isolated from single-cell suspensions using a magnetic negative selection kit (Miltenyi Biotec, #130-104-075), following the manufacturer’s protocol. Isolation efficiency was validated by flow cytometry (for FACS panel, see Supplementary Table 1). Splenocytes were activated by addition of the TLR7 agonist R848 (Invivogen, #tlrl-r848-5) at a concentration of 25 ng/mL at the same time as HMPSNE.

### Generation of bone marrow-derived dendritic cells (BMDC)

BMDC were generated from 6–16-week-old female C57BL/6J mice as described [22]. Bone marrow was flushed from femurs and tibiae with PBS and treated with Pharm Lyse buffer. After filtration through a 40 μm strainer, cells were seeded in 6-well plates at 2.5×10⁶ cells/mL in complete RPMI medium with 20 ng/mL GM-CSF (Peprotech, #315-03) and cultured in a 37°C, 5% CO_2_ humidified environment. The medium was refreshed on days 2 and 3 after isolation. On day 6, BMDC were harvested and plated into 96-well plates at 1-1.5×10⁵ cells/well. Purity was verified by flow cytometry (for FACS panel, see Supplementary Table 1).

### BrdU incorporation assay

To assess proliferation, 1.5×10⁴ Renca-H2-K^b^ GFP^+^ cells were seeded per well in 96-well plates and allowed to adhere overnight. Cells were then treated with varying concentrations of HMPSNE and labeled with 10 μM bromodeoxyuridine (BrdU) for 24 h. BrdU incorporation was analyzed by ELISA (Roche, #11647229001), following the manufacturer’s instructions. Absorbance at 450 nm was measured using a Clariostar Plus plate reader (BMG Labtech).

### Metabolic cell viability assay

Splenocytes and BMDC (100,000 cells/well) were seeded in 96-well plates and treated with HMPSNE immediately after plating. Renca-H2-Kb GFP+ cells (5,000 cells/well) were seeded and allowed to adhere for 24 h prior to HMPSNE treatment. Metabolic activity was measured by CellTiter-Glo® (Promega, #G7572) according to the manufacturer’s instructions using a Clariostar Plus plate reader.

### Flow cytometry

Flow cytometry was performed using a NovoCyte 3000 (ACEA Biosciences). Single-cell suspensions from tissues or cultures were stained with Zombie Violet™ (BioLegend, #423114) to exclude dead cells. Cells were incubated with anti-CD16/32 (BioLegend, #101320) and fluorochrome-conjugated antibodies (Supplementary Table 1) for 30 min at 4 °C. Staining was performed in FACS buffer (PBS + 0.5% BSA + 2% EDTA). Apoptosis was assessed using Annexin V-Pacific Blue (BioLegend, #640917) and propidium iodide (Sigma, #P4170) per protocol. Data were analyzed using FlowJo™ v10.10.0 (FlowJo LLC).

### CD8^+^ T-cell proliferation assay

CD8^+^ T cells isolated from OT-I mice were resuspended at 10×10⁶ cells/mL in PBS with 0.5 μM CFSE (BioLegend, #423801) for 10 min at room temperature. Cells were plated in complete RPMI medium supplemented with 50 μM 2-mercaptoethanol (Gibco, #31350010) at 100,000 cells/well in 96-well plates. T cells were activated with mouse IL-2 (30 U/mL; Gibco, #210-12-10UG) and CD3/CD28 Dynabeads (1:1 bead-to-cell ratio, Gibco, #1145D) during exposure to HMPSNE. After 24 and 48 h, beads were removed using a DynaMag magnet (Invitrogen), and cells were analyzed by flow cytometry (Supplementary Table 1).

### CD8^+^ T cell-mediated anti-tumor cytotoxicity

Renca-H2-Kb GFP^+^ cells (1,000 cells/well) were seeded in 96-well plates, allowed to adhere for 24 h, then treated with HMPSNE. Tumor cells were pulsed with 2 μg/mL SIINFEKL peptide (InvivoGen, #vac-sin) for 4 h at 37 °C. OT-I CD8^+^ T cells (5,000 cells/well) were added. Tumor cell proliferation was monitored by GFP intensity using the Incucyte® Live-Cell Analysis System (Sartorius). Anti-PD-L1 antibody (10 μg/mL; InvivoGen, #pdl1-mab15-1) was added where indicated.

### Transcriptomic analysis

Gene expression analysis and survival correlation from the TCGA Kidney Clear Cell Carcinoma (KIRC) cohort (n= 533; https://www.cancer.gov/tcga) were performed using the UCSC Xena Browser [23] (https://xenabrowser.net). Patients were dichotomized into high and low expression groups based on the median cutoff values. Survival differences were evaluated using the log-rank test calculated by the built-in statistical pipeline. To ensure consistency, only primary tumor samples (TCGA sample type code: 01) were retained for analysis. Duplicate entries and rare sample types, such as “additional new primary” tumors, were removed according to the UCSC Xena Browser’s recommended preprocessing guidelines. Figures were exported directly from the platform. The direct links to the Kaplan-Meier curves are provided below:

CBS: https://xenabrowser.net/?bookmark=846161bebc4b7b607a75161982f5001c

CSE/CTH: https://xenabrowser.net/?bookmark=9a2b100681a240ec66802ba2f6f2e10c

3-MST/MPST: https://xenabrowser.net/?bookmark=b32c2f86318d5acab2088193317fac61

### Immunohistochemistry (IHC) analysis of H₂S-synthesizing enzymes in normal kidney and RCC

IHC data were obtained from the Human Protein Atlas (HPA, version 24; https://v24.proteinatlas.org) [24]. Images and expression annotations for both normal kidney and RCC samples were retrieved from the “Tissue” and “Pathology” sections of the HPA. Normal kidney data were used to describe baseline distribution of each enzyme in specific nephron segments and glomerular structures, while RCC data were analyzed for presence or absence of staining, staining intensity (not detected, low, medium, high), and proportion of tumor cells stained, as provided by the HPA (CBS, 3-MST: n=12; CSE/CTH: n=11). Representative high-resolution images were downloaded for CBS (https://images.proteinatlas.org/1223/5627_A_9_5.jpg), CSE/CTH (https://images.proteinatlas.org/23300/51662_A_7_6.jpg), and 3-MST (https://images.proteinatlas.org/1240/4178_A_7_2.jpg) for presentation in Figure 1.

**Figure 1.**
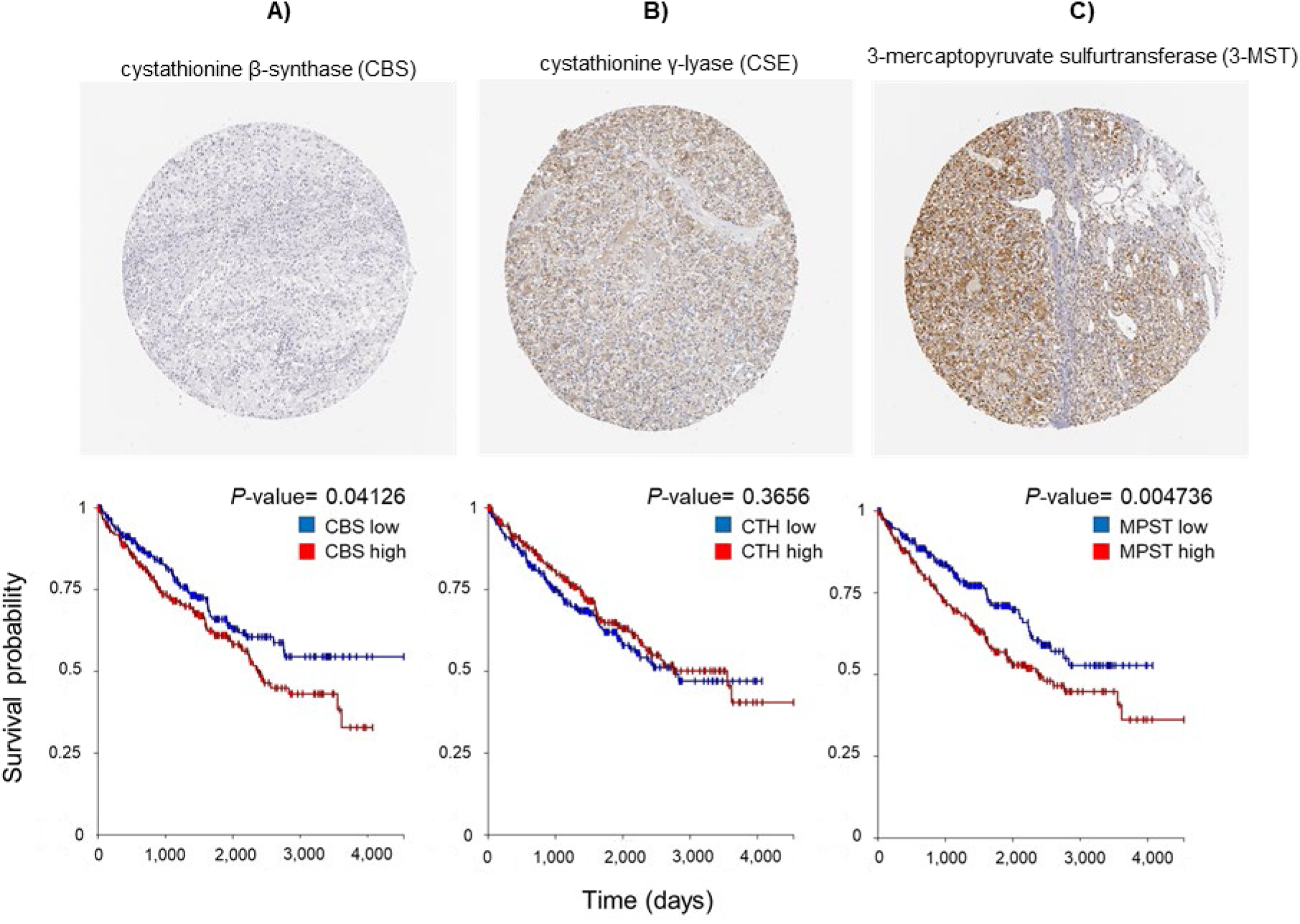
Expression of H_2_S-synthesizing enzymes and correlation with survival in human RCC. **Top row:** Representative images of immunohistochemistry (IHC) staining of human renal cell adenocarcinoma (RCC) samples from the Human Protein Atlas showing expression of cystathionine β-synthase (CBS), cystathionine γ-lyase (CSE), and 3-mercaptopyruvate sulfurtransferase (3-MST). **Bottom row:** Kaplan–Meier curves of ccRCC patients stratified by high vs. low mRNA expression levels of CBS, CTH (encoding CSE), and MPST (encoding 3-MST). Patients were dichotomized into high vs. low gene expression groups based on the median expression values as threshold. Data derived from The Cancer Genome Atlas (TCGA), KIRC cohort (n=533; low-expression group: n=267, high-expression group: n=266).

### Statistical analysis

Statistical analyses were performed using GraphPad Prism. In vitro data are shown as mean ± SD of biological replicates unless indicated otherwise. Statistical significance was assessed using two-way ANOVA with Dunnett’s or Tukey’s multiple comparison tests as indicated. A p-value ≤ 0.05 was considered significant.

### Ethics statement

No new human samples were collected for this study, and all analyses for human tissues were performed on de-identified, previously published datasets.

## Results

### 3-MST is highly expressed in human renal cell cancer and correlates with poor prognosis

To assess the expression of H₂S-synthesizing enzymes in human renal cell cancer (RCC), we examined immunohistochemistry (IHC) images from the Human Protein Atlas (HPA) [25]. In normal kidney, cystathionine β-synthase (CBS) was not detectable, whereas cystathionine γ-lyase (CSE) was expressed at high levels in proximal tubules and at lower levels in distal tubules but was not detectable in glomeruli. The staining pattern for 3-mercaptopyruvate sulfurtransferase (3-MST) showed strong positivity in the epithelial cells of the proximal tubules, whereas the distal tubules showed weak staining. Vascular structures were negative. In the glomerulus, the parietal epithelial cells of the Bowman’s capsule were strongly positive. In renal cell adenocarcinoma, which arises from cells of the proximal tubule, CBS was not detected in any of the HPA samples (n=12) (representative image in Figure 1A, top row). CSE was not detected in 6 of 11 samples and detected at low (4/11) or medium (1/11) levels in the others (Figure 1B, top row). In contrast, 3-MST was present at low (6/12) to medium (2/12) levels in most cases, with clear staining in over 25% of tumor cells in the majority of analyzed samples (Figure 1C, top row).

We next examined the association between expression of H₂S-synthesizing enzymes and patient survival using mRNA expression data from The Cancer Genome Atlas (TCGA) KIRC cohort (n=533). Patients were stratified into high and low expressers based on median transcript levels of *MPST* (encoding 3-MST), *CBS*, or *CTH* (encoding CSE). Kaplan-Meier survival analysis showed that low *CBS* expression was associated with a non-significant trend toward improved survival (Figure 1A, bottom row), and that no difference was observed between high and low *CTH* expression groups (Figure 1B, bottom row). In contrast, low *MPST* expression was significantly associated with improved overall survival (p < 0.004), suggesting a potential role for 3-MST in promoting tumor aggressiveness (Figure 1C, bottom row). These data suggest that among the three major H₂S-producing enzymes, 3-MST is the most highly expressed in human RCC and that its elevated expression may correlate with worse clinical outcomes. To explore the functional relevance of 3-MST in renal cancer, we next investigated the effects of its pharmacological inhibition in renal cancer cells.

### Pharmacological inhibition of 3-MST reduces intracellular H₂S in renal cancer cells

First, we examined the impact of 3-MST inhibition on endogenous H₂S production in renal cancer cells by quantifying intracellular H₂S levels in murine Renca renal carcinoma cells treated with the selective 3-MST inhibitor HMPSNE. Fluorescent detection using the H₂S-sensitive AzMC probe revealed a clear signal in untreated cells, which decreased upon treatment with increasing concentrations of HMPSNE (Figure 2A). Quantification of the AzMC signal confirmed a dose-dependent reduction in fluorescence, indicating reduced intracellular H₂S levels (Figure 2B). This decrease was statistically significant at all tested concentrations of the inhibitor. These results confirm that HMPSNE effectively suppresses H₂S production in Renca cells and validate its utility for functional studies of 3-MST inhibition in renal cancer cells.

**Figure 2.**
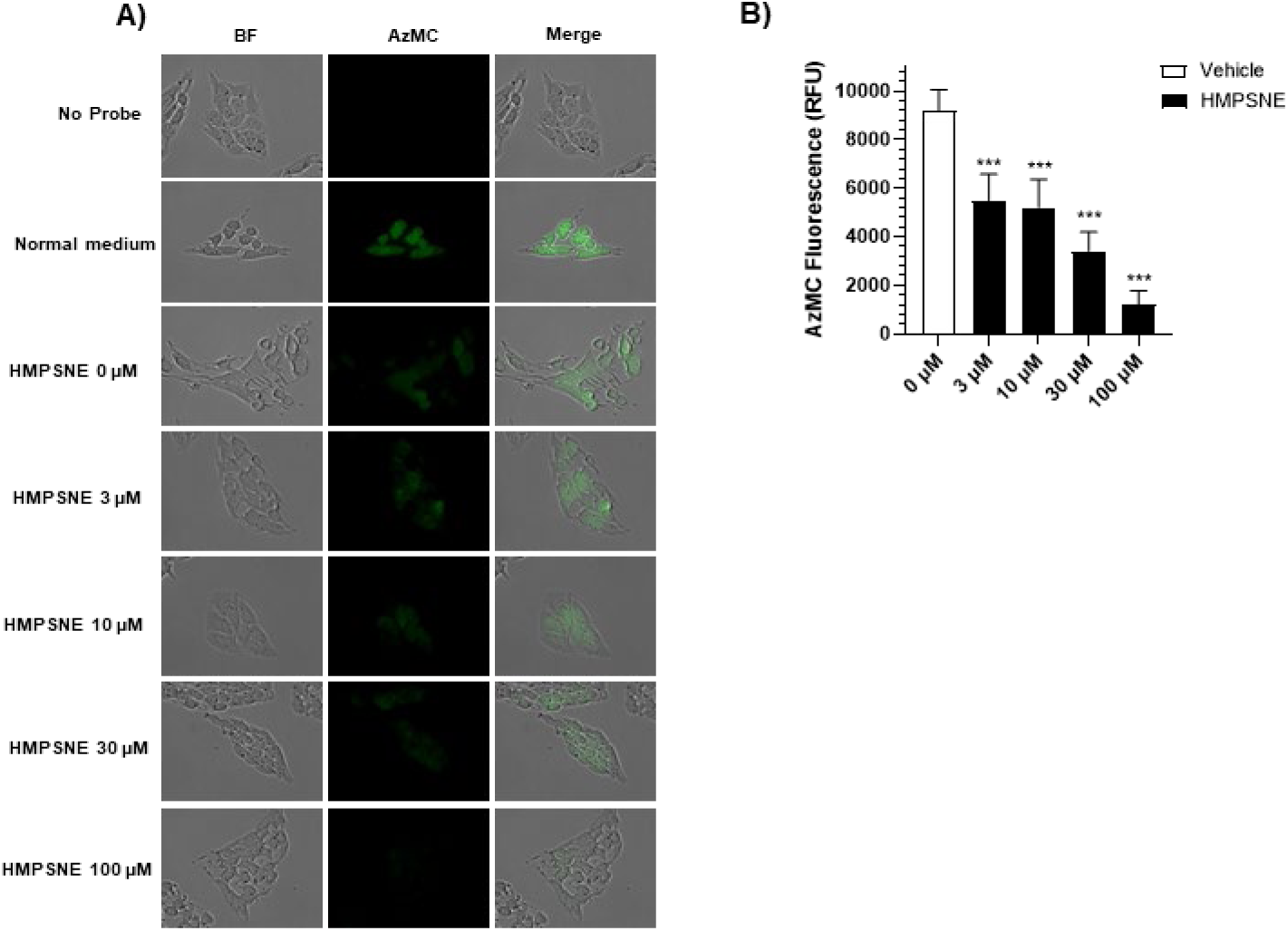
H₂S production in renal cancer cells following pharmacological inhibition of 3-MST. Cellular H₂S content in Renca cells was quantified using the AzMC (7-azido-4-methylcoumarin) probe. (A) Representative bright-field (BF) and AzMC fluorescence images of cells treated with increasing concentrations of HMPSNE for 48 h. (B) Quantification of AzMC fluorescence signal across HMPSNE concentrations. Data represent mean ± SD from six independent experiments. Statistical analysis was performed using ordinary one-way ANOVA. ***p < 0.001 versus vehicle control (0 µM HMPSNE).

### 3-MST inhibition reduces proliferation and induces apoptosis in Renca cells

To investigate the functional impact of 3-MST inhibition on tumor cell proliferation, GFP-labeled Renca cells (Renca H2-Kb GFP) were treated with increasing concentrations of HMPSNE. Live-cell imaging revealed a concentration-dependent reduction in proliferation (Figures 3A and C). Consistently, a reduction in metabolic activity was observed from 50 µM HMPSNE at 48 hours post-treatment (Figure 3B). The antiproliferative effect was further confirmed by BrdU incorporation, which showed reduced DNA synthesis at the highest inhibitor concentration after 48 hours (Figure 3D). Notably, the decrease in proliferation was accompanied by a dose-dependent increase in apoptotic cells at both 24 and 48 hours after treatment (Figure 3E).

**Figure 3.**
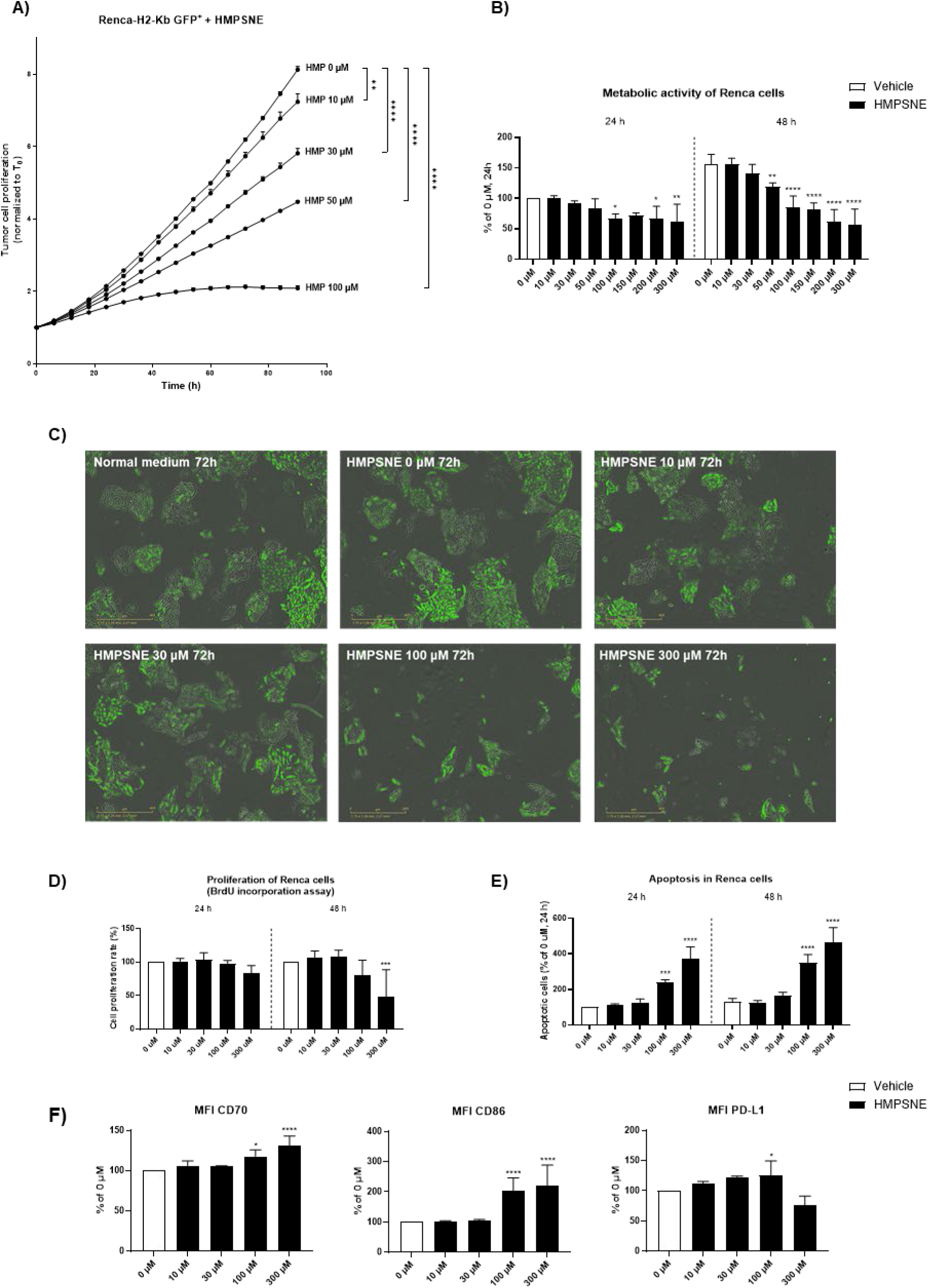
Effect of 3-MST inhibition on viability and phenotype of renal cancer cells. Renca-H2-Kb GFP⁺ cells were treated with increasing concentrations of HMPSNE. (A) Live-cell imaging expressed as GFP signal normalized to the starting point (T₀ = 1) for each well. Data represent mean ± S.E.M. from one representative experiment out of three with three technical replicates. (B) Metabolic activity measured by CellTiter-Glo® assay at 24 h and 48 h. (C) Representative images of Renca-H2-Kb GFP⁺ cells treated with increasing concentrations of HMPSNE for 72 h. Images were acquired using the Incucyte® Live-Cell Analysis System (20× objective). (D) Cell proliferation determined by BrdU incorporation. (E) Annexin V-positive, PI-negative apoptotic cells measured by flow cytometry. (F) Expression of CD70, CD86, and PD-L1 (mean fluorescence intensity, MFI) on Renca-H2-Kb GFP⁺ cells determined by flow cytometry. Data represent mean ± SD from three independent experiments. Statistical analysis was performed using two-way ANOVA with Tukey’s post hoc test for (A) and Dunnett’s post hoc test for (B, D-E). ****p < 0.0001; ***p < 0.001; **p < 0.01; *p < 0.05 versus vehicle control (0 µM HMPSNE).

In addition to reducing tumor cell viability, HMPSNE altered the immunogenic phenotype of Renca cells. Flow cytometry analysis showed increased surface expression of CD70 and CD86, both co-stimulatory molecules involved in T cell activation [26,27], at 100 and 300 µM HMPSNE, and of PD-L1, an immune checkpoint ligand, at 100 µM HMPSNE (Figure 3F). No significant changes were observed in the expression of CD155, Galectin-9, or MHC class I (data not shown). These findings demonstrate that pharmacological inhibition of 3-MST suppresses Renca cell proliferation, promotes their apoptosis, and enhances the expression of immunostimulatory and immune checkpoint molecules, potentially increasing tumor cell visibility to the immune system. Given these immunomodulatory effects on tumor cells, we next examined whether 3-MST inhibition also directly influences immune cell function.

### HMPSNE has limited cytotoxicity on immune cells and supports T-cell activation

To evaluate the effects of 3-MST inhibition on the viability and functional activation of immune cells, mouse splenocytes were treated with increasing concentrations of HMPSNE. Murine splenocytes are a heterogeneous population of immune cells composed of B and T lymphocytes as well as myeloid cells such as monocytes and dendritic cells [28]. A trend towards a reduction in the metabolic activity of total splenocytes was observed at 24 and 48 hours at high concentrations of HMPSNE (Figure 4A, left panel). 3-MST inhibition during splenocyte activation with a TLR7 agonist, R848, showed a significant decrease in metabolic activity only at the highest concentration of HMPSNE, 300 µM, after 48 hours (Figure 4A, right panel). Furthermore, HMPSNE enhanced T cell activation within splenocytes activated by R848, as measured by surface expression of the early activation marker CD69. Although no change was seen on total activated splenocytes (defined as CD45-positive cells), a significant increase in CD69 mean fluorescence intensity was observed in total CD3⁺ T cells at 300 µM, in CD4⁺ helper T cells at both 100 and 300 µM, and in CD8⁺ cytotoxic T cells at 300 µM (Figure 4B).

**Figure 4.**
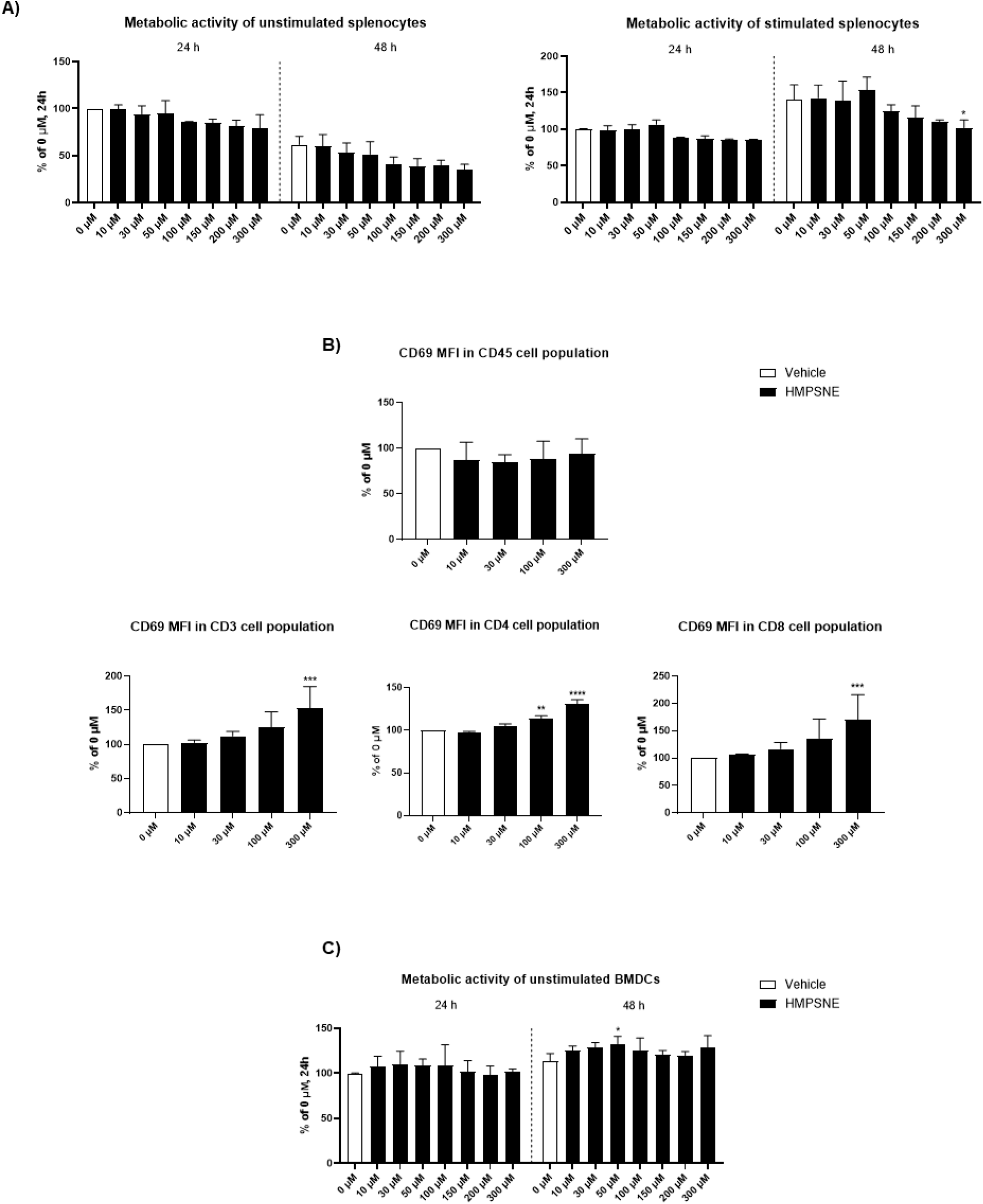
Effect of 3-MST inhibition on viability and activation of immune cells. (A) Mouse splenocytes were either left unstimulated or stimulated with R848 and treated at the same time with increasing concentrations of HMPSNE. Cell viability assessed at 24 h and 48 h using the CellTiter-Glo® luminescent assay, expressed as percentage of the unstimulated control (0 µM HMPSNE). (B) Flow cytometric analysis of CD69 expression on R848-activated CD45⁺ splenocytes, total T cells (CD3⁺), and T-cell subtypes: helper (CD4⁺) and cytotoxic (CD8⁺), presented as percentage of the control MFI. Gating was performed sequentially on CD45⁺ cells, CD3⁺ T cells, and respective subpopulations. (C) Metabolic activity was assessed at 24 h and 48 h of the unstimulated bone marrow-derived dendritic cells (BMDC) using the CellTiter-Glo® (CTG) luminescent assay, with values expressed as percentage of the control (0 µM HMPSNE). Data represent mean ± SD from three independent experiments. Statistical analysis was performed using two-way ANOVA with Dunnett’s post hoc test. ****p < 0.0001; ***p < 0.001; **p < 0.01; *p < 0.05 versus control (0 µM HMPSNE).

The effect of 3-MST inhibition on dendritic cells was also examined. Dendritic cells are antigen-presenting cells that play an essential role in the initiation and orchestration of antitumor immune responses [29]. Dendritic cells were differentiated from mouse bone marrow (bone marrow-derived dendritic cells, BMDC) and exposed to HMPSNE for up to 48 hours. No decrease in metabolic activity of BMDC was observed, even at the highest concentrations of the inhibitor (Figure 4C). These data suggest that while high concentrations of HMPSNE may modestly affect viability of some immune cell subtypes, the compound also enhances T-cell activation in both the CD4⁺ and CD8⁺ subsets. These findings suggest that 3-MST inhibition may enhance T-cell activation, supporting a potential role in modulating antitumor immune responses.

### 3-MST inhibition impairs proliferation of CD8⁺ T cells at high concentrations

To assess how inhibition of 3-MST influences CD8⁺ T cell proliferation and phenotype, purified CD8⁺ T cells were labeled with CFSE and stimulated with CD3/CD28-coated beads and IL-2 in the presence of HMPSNE. Flow cytometric analysis of CFSE dilution revealed that proliferation was unaffected at lower concentrations of HMPSNE (10–30 µM), but reduced at 100 and 300 µM HMPSNE, as evidenced by fewer CFSE_low_ cells (Figure 5A–B). This inhibitory effect extended to all examined CD8⁺ T cell subsets, including naïve (CD62L^hi^/CD44^lo^), central memory (CD62L^hi^/CD44^hi^) and effector memory (CD62L^lo^/CD44^hi^) populations, which showed decreased expansion at higher HMPSNE concentrations (Figure 5C). Despite the reduced proliferation, expression of the activation markers PD-1 and CD25 was maintained, suggesting that activation status was at least partially preserved (Figure 5D). These results indicate that while high-dose HMPSNE can impair CD8⁺ T cell proliferation, it does not suppress activation, highlighting a concentration-dependent effect of 3-MST inhibition on T cell function.

**Figure 5.**
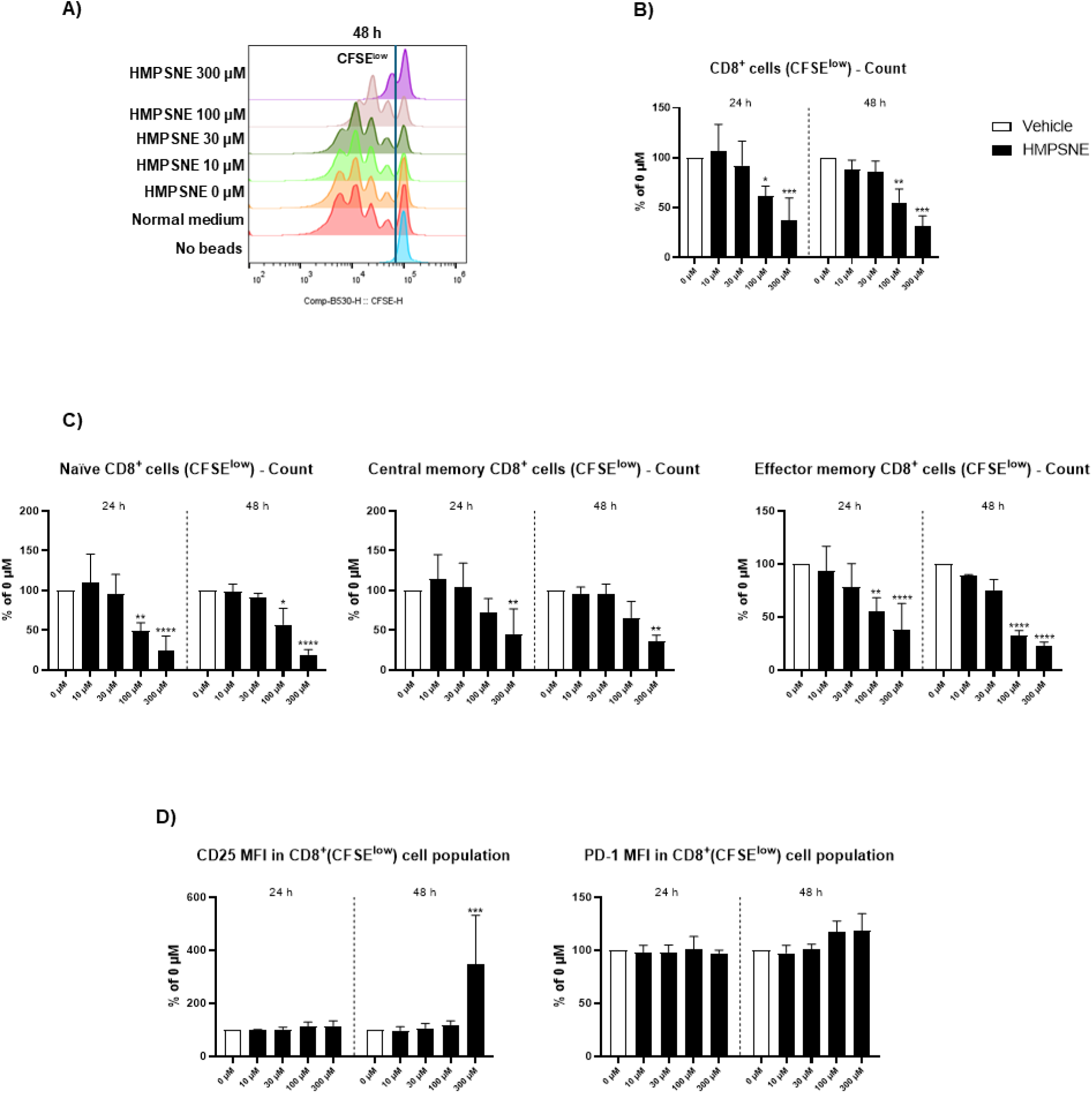
Effect of HMPSNE on phenotype and proliferation of CD8⁺ T cells. CD8⁺ T cells were labeled with CFSE, activated with Dynabeads (CD3/CD28) and IL-2 and treated with increasing concentrations of HMPSNE. (A) Representative CFSE histograms showing cell division peaks. (B) Quantification of CFSE^low^ CD8⁺ T-cells, expressed as percentage of control (0 µM HMPSNE). (C) Quantification of CFSE^low^ CD8⁺ T-cell subsets, including naïve (CD62L^hi^/CD44^lo^), central memory (CD62L^hi^/CD44^hi^), and effector memory (CD62L^lo^/CD44^hi^) populations, expressed as percentage of control (0 µM HMPSNE). (D) PD-1 and CD25 expression on CFSE^low^ CD8⁺ T cells, presented as percentage of control MFI. Data represent mean ± SD from three independent experiments. Statistical analysis was performed using two-way ANOVA with Dunnett’s post hoc test. ****p < 0.0001; ***p < 0.001; **p < 0.01; *p < 0.05 versus control (0 µM HMPSNE).

### 3-MST inhibition enhances CD8⁺ T cell-mediated cytotoxicity and cooperates with PD-L1 blockade

To evaluate whether 3-MST inhibition affects antigen-specific T cell-mediated killing of tumor cells, we co-cultured SIINFEKL-loaded Renca-H2-K^b^ GFP⁺ target cells with CD8⁺ T cells isolated from OT-I mice in the presence or absence of HMPSNE. SIINFEKL is an immunodominant peptide epitope from ovalbumin that is specifically recognized by CD8⁺ T cells from OT-I transgenic mice. GFP-based proliferation tracking revealed that growth of tumor cells was reduced in the presence of OT-I cells when they were loaded with SIINFEKL, and that this effect was further potentiated by HMPSNE in a dose-dependent fashion (Figure 6A). In the absence of SIINFEKL loading, OT-I cells exerted only a modest inhibitory effect on tumor cell growth, consistent with limited bystander killing (data not shown). These results demonstrate that pharmacological inhibition of 3-MST enhances antigen-specific CD8⁺ T cell– mediated killing of tumor cells in a dose-dependent manner.

**Figure 6.**
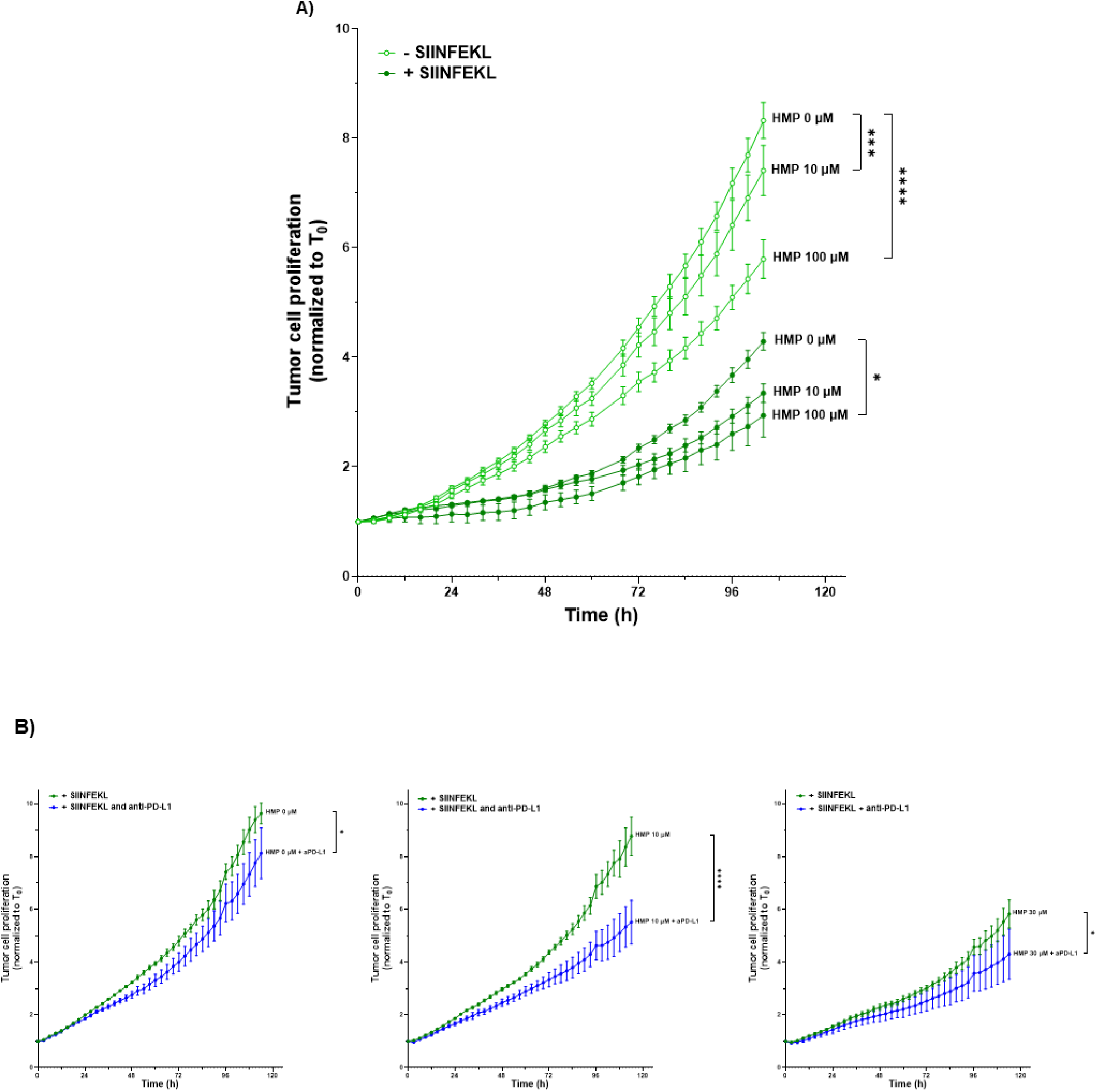
Impact of pharmacological inhibition of 3-MST on CD8⁺ T cell-mediated cytotoxicity. CD8⁺ T cell-mediated cytotoxicity was assessed using Renca-H2-Kb GFP⁺ tumor cells either loaded with SIINFEKL or left unloaded, and co-cultured with CD8⁺ T cells isolated from OT-1 mice in the presence of increasing concentrations of HMPSNE. (A) Proliferation curves of tumor cells based on GFP signal, normalized to the starting point (T₀ = 1) for each well. (B) Tumor cell proliferation over 114 h with increasing concentrations of HMPSNE and/or anti-PD-L1, expressed as values normalized to T₀. Data represent mean ± SEM from one representative experiment out of three with three technical replicates. Statistical analysis was performed using two-way ANOVA with Tukey’s post hoc test. ****p < 0.0001; ***p < 0.001; *p < 0.05 versus indicated sample.

We next tested whether immune checkpoint inhibition further amplifies T cell-mediated cytotoxicity. Addition of anti-PD-L1 antibody significantly enhanced OT-I-mediated cytotoxicity against SIINFEKL-loaded targets (Figure 6B, left panel). Importantly, combining anti-PD-L1 with partial inhibition of 3-MST activity –– achieved by low concentrations of HMPSNE (10–30 µM, Figure 6B, middle and right panels) –– further increased tumor cell suppression compared to checkpoint inhibition in the absence of HMPSNE. These findings suggest that 3-MST inhibition can enhance CD8⁺ T cell effector function and sensitize tumor cells to immune checkpoint blockade, potentially by increasing their immunogenicity and/or altering redox-dependent survival pathways.

## Discussion

H₂S is recognized as a pivotal gasotransmitter in redox biology, exerting either cytoprotective or cytotoxic effects depending on its concentration, cellular context, and disease state [30–32]. Within the tumor microenvironment, dysregulated H₂S metabolism is increasingly recognized as a driver of malignant progression, metabolic reprogramming, and immune escape [33]. Among the three main H₂S-synthesizing enzymes, CBS, CSE, and 3-MST, our analysis of publicly available datasets identified 3-MST as the most abundantly expressed at the protein level in both healthy kidney tissue and RCC. This pattern is consistent with previous reports of elevated 3-MST in other malignancies, including colon adenocarcinoma, lung adenocarcinoma, and urothelial carcinoma [14]. We further show that high 3-MST expression is associated with reduced survival in RCC patients, implicating this enzyme and its product, H₂S, in disease progression.

Pharmacological inhibition of 3-MST with HMPSNE led to a dose-dependent reduction in intracellular H₂S levels in Renca cells, consistent with prior reports of HMPSNE specificity [21]. This inhibition suppressed proliferation and induced apoptosis, reinforcing earlier observations that endogenous H₂S protects cancer cells from oxidative stress and apoptosis through multiple mechanisms [8,34]. Notably, 3-MST blockade also triggered immunogenic remodeling of tumor cells, with increased expression of co-stimulatory molecules (CD70, CD86) and the immune checkpoint PD-L1. These changes mirror redox-driven immune phenotypes described previously, where reactive oxygen species (ROS) enhance antigen presentation machinery and checkpoint expression [35,36]. This immunogenic shift may render tumor cells more susceptible to immune recognition and therapeutic intervention, especially in the context of checkpoint blockade.

A striking aspect of our findings is the differential sensitivity of tumor versus immune cells to 3-MST inhibition. Whereas Renca cells showed a marked loss of viability, splenocytes and dendritic cells were comparatively resistant to the cytotoxic effects of HMPSNE. Furthermore, HMPSNE did not suppress—and in fact enhanced—lymphocyte activation, as shown by increased CD69 expression on both CD4⁺ and CD8⁺ T cells following Toll-like receptor (TLR) stimulation. CD69, an early activation marker, can be induced via TLR or cytokine signals independent of antigen-specific T cell receptor engagement [29]. This suggests that modulating the reactive sulfur homeostasis via 3-MST inhibition may preferentially promote effector immune pathways over regulatory or tolerogenic programs. However, at higher HMPSNE concentrations, CD8⁺ T cell proliferation was impaired despite preserved activation marker expression, suggesting a dose-dependent threshold in redox homeostasis—a concept consistent with the role of H₂S as a redox rheostat, where both depletion and excess can compromise immune function [5,37].

The most compelling translational insight from this study is the cooperative effect between 3-MST inhibition and CD8⁺ T cell-mediated cytotoxicity. Pharmacological inhibition of 3-MST promoted the cytotoxic activity of CD8^+^ T cells against antigen-specific tumor targets and further enhanced efficacy when combined with PD-L1 blockade. This cooperative effect suggests that H₂S-targeting metabolic interventions can sensitize cancer cells to immune attack, possibly by boosting immune costimulatory signals, altering redox-sensitive survival pathways, or reducing metabolic competition at the tumor–immune interface. While further studies are needed to validate these findings using in vivo preclinical studies and human RCC models, our data support a model in which 3-MST sustains a tumor-permissive redox state that limits immune activation and promotes tumor cell survival. Targeting this pathway may represent a therapeutic strategy to simultaneously disrupt tumor metabolism and reawaken antitumor immunity.

## Acknowledgements

This work was supported by the Swiss National Science Foundation (Projects 182317 and IZSTZ0_198887), the Swiss Cancer Research Foundation KFS 4535-08-2018-R and the Novartis Foundation.

---

**Supplementary Table 1.**
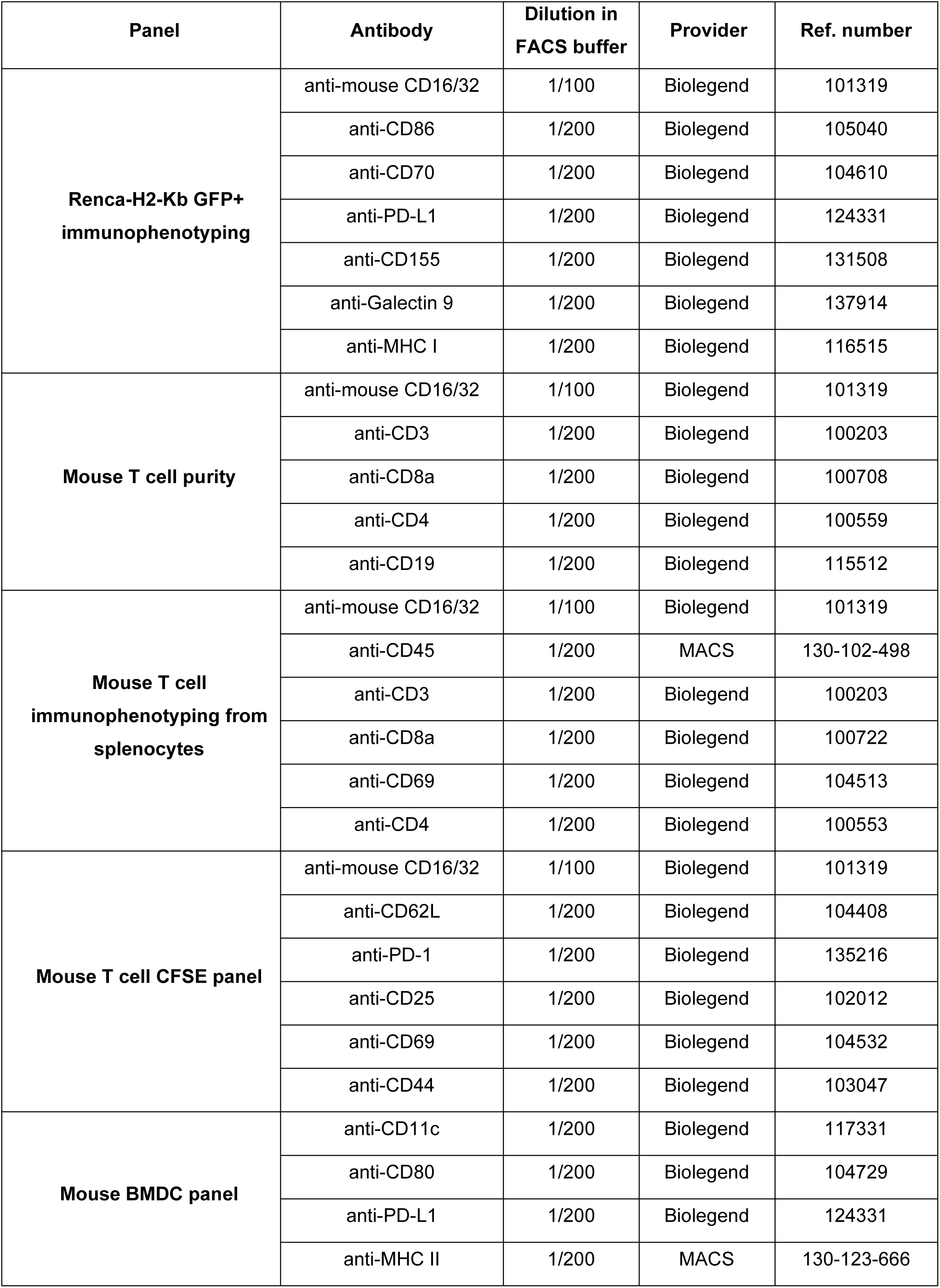
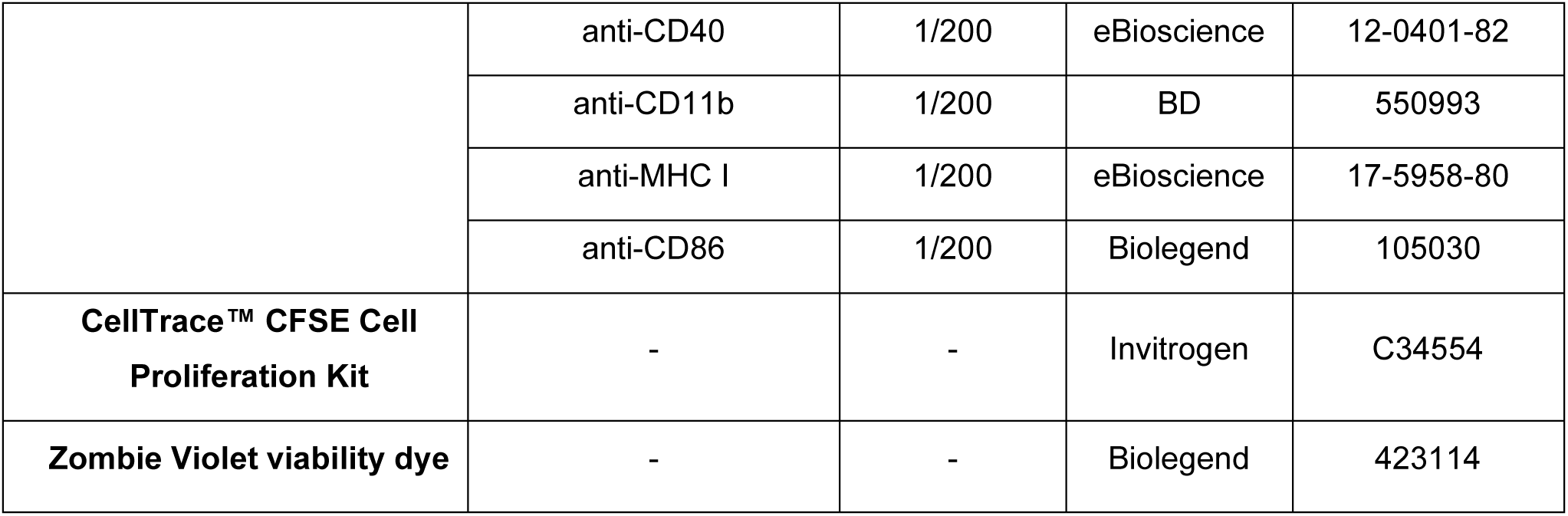
FACS panels and amine-reactive dyes.

